# Plasma-Polymerized Polyurethane/Reduced Graphene Oxide Electrospun Patch for Cardiac Tissue Engineering

**DOI:** 10.64898/2026.11.22.768432

**Authors:** Clotilde Jugie, Ali Mousavi, Alireza Nemati, Michael Wertheimer, Abdellah Ajji, Houman Savoji

**Author notes:** These authors contributed equally to the work (Clotilde Jugie, Ali Mousavi). Corresponding Author: Houman Savoji - Institute of Biomedical Engineering, Polytechnique Montréal, Montréal, Québec, Canada.

## Abstract

Cardiovascular diseases remain the leading cause of mortality worldwide. Traditional treatments, including heart transplantation, are limited by the scarcity of donor organs and the need for lifelong immunosuppression, highlighting the urgent need for alternative therapies. Cardiac patches represent a promising approach to restoring heart function by providing structural support and promoting cardiac tissue regeneration. Here, we developed biocompatible and conductive electrospun cardiac patches, aiming to mimic the structural, mechanical, and electrical properties of the native myocardium. polyurethane (PU) was selected as the base material due to its favorable mechanical properties and biocompatibility. Reduced graphene oxide (rGO) was incorporated into the electrospun PU fibers to enhance electrical conductivity, and a low-pressure plasma-polymerized nitrogen-rich (LPPE:N) coating was applied to improve cell adhesion. The fabricated patches were characterized through various techniques, including scanning electron microscopy, X-ray photoelectron spectroscopy, tensile testing, and conductivity measurements. Also, *in vitro* cytotoxicity assay using primary cardiac fibroblasts confirmed the biocompatibility of the patches. These physicochemical, electromechanical, and biological characterizations suggest that the rGO/LPPE:N-functionalized PU patches demonstrate promising potential for cardiac tissue engineering applications.

## 1. Introduction

Cardiovascular diseases (CVDs) represent the foremost cause of mortality worldwide, contributing to 32% of global deaths and imposing a substantial health burden [1]. Among these, ischemic heart diseases (IHDs), primarily driven by atherosclerosis-induced coronary artery blockages, account for 16% of global deaths [2]. The resulting damage to the myocardium involves irreversible cardiomyocyte loss and scar tissue formation, leading to progressive cardiac dysfunction and, in severe cases, cardiac arrest [3, 4]. While heart transplantation remains a viable treatment option, its limitations, such as donor shortages, prolonged waiting times, immunosuppression requirements, and surgical risks, highlight the urgent need for alternative therapeutic approaches [5]. Emerging options like cell therapy and xenotransplantation show potential but face significant challenges, including low cell retention, inconsistent results, and immune complications [6, 7].

Cardiac tissue engineering offers a promising alternative, employing advanced techniques like electrospinning, 3D bioprinting, and hydrogel-based scaffolds to replicate the structural and functional attributes of native cardiac tissue [8–10]. Electrospinning, in particular, excels in creating nanofibrous scaffolds that mimic the extracellular matrix (ECM), enabling enhanced cell adhesion, proliferation, and integration [11, 12]. By tailoring electrospun fiber properties, such as mechanical strength, elasticity, and electrical conductivity, researchers aim to fabricate cardiac patches that can restore myocardial functionality [13, 14].

For effective cardiac repair, scaffolds must closely replicate the mechanical, structural, and electrical properties of the myocardium [15, 16]. Mechanically, the myocardium exhibits a Young’s modulus of 10–20 kPa during systole and 200–500 kPa during diastole, along with tensile strength ranging between 3 and 15 kPa [13, 17]. The myocardium also undergoes 20-30% mechanical strain during the cardiac cycle, highlighting the dynamic environment that cardiac patches must endure [18, 19]. Structurally, the native ECM comprises nanoscale collagen fibers (50-500 nm in diameter) arranged to support cell organization and stress distribution, and scaffolds should reflect this alignment for optimal integration [20, 21]. Electrically, the myocardium demonstrates longitudinal conductivity of approximately 2.1 mS/cm and transverse conductivity of 0.5 mS/cm, critical for synchronized contractions and effective blood pumping [22, 23].

Materials play a pivotal role in achieving these properties. Synthetic polymers like polyurethane (PU) and polycaprolactone (PCL) offer customizable mechanical characteristics, while natural polymers such as collagen and gelatin enhance biocompatibility [24, 25]. Conductive materials, including reduced graphene oxide (rGO) and carbon nanotubes (CNTs), further improve electrical functionality, enabling scaffolds to facilitate cardiac signaling [26, 27]. To enhance cell adhesion and integration, bioactive coatings such as plasma-polymerized films introduce functional groups (e.g., amines and carboxyl groups) that promote cellular interactions and scaffold integration [28–30].

This research aims to fabricate electrospun cardiac patches that replicate the myocardium’s mechanical, structural, and electrical properties. Medical-grade PU was selected for its elastomeric behavior, and rGO was incorporated to enhance conductivity. The design features a combination of aligned and randomly oriented fibers to provide optimal mechanical performance and support cell alignment. A low-pressure plasma-polymerized nitrogen-rich (LPPE:N) coating was also applied to improve cell adhesion and integration. **Fig. 1** represents a schematic of the developed patch and an overview of the study, which aims to advance bioengineered solutions for effective myocardial regeneration.

**Fig. 1.**
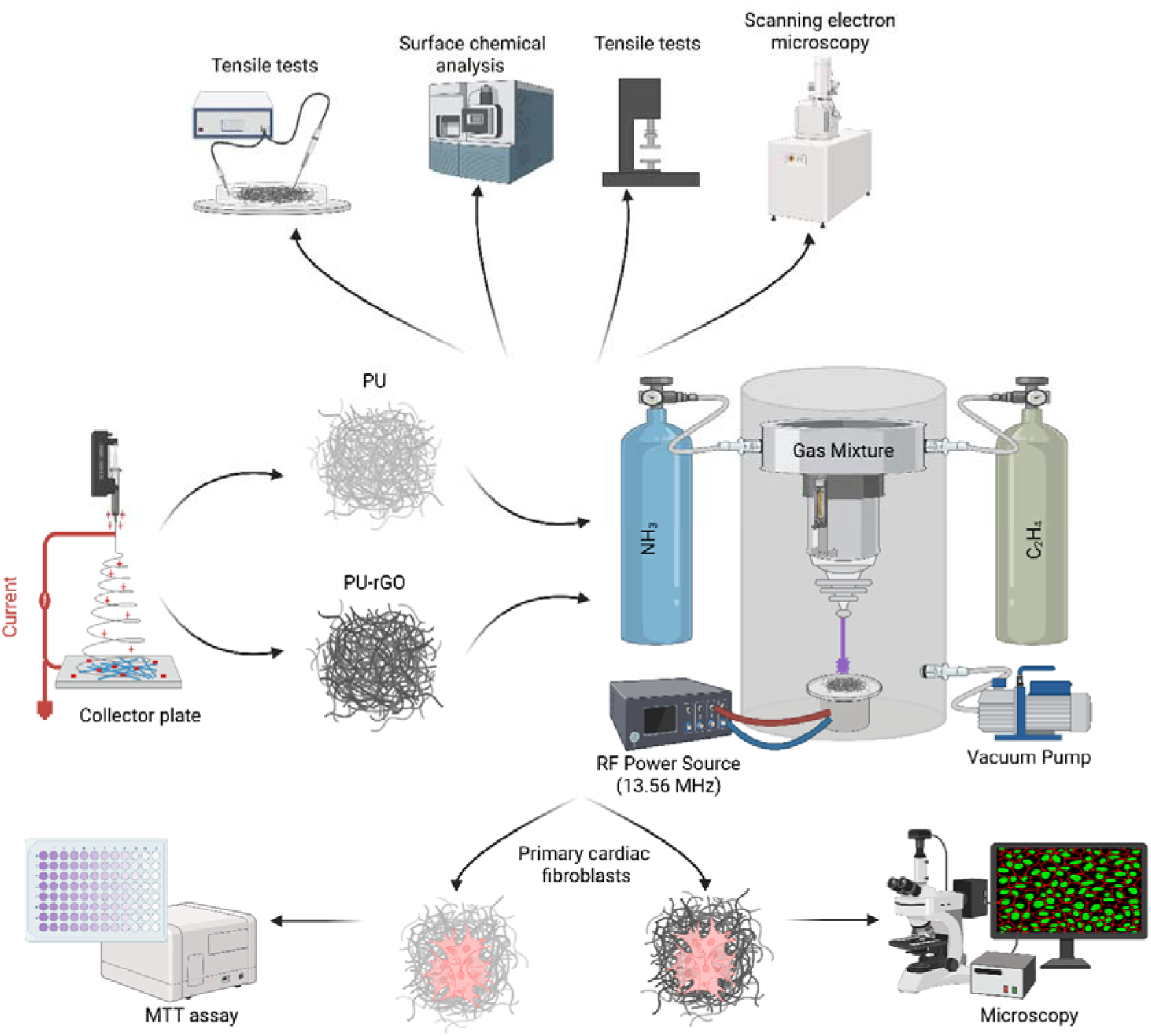
Schematic overview of the fabrication, surface functionalization, characterization, and biological evaluation of electrospun PU and PU-rGO fibrous mats. PU and PU-rGO fibers were produced by electrospinning and functionalized using LPPE:N. The materials were characterized by SEM, surface chemical analysis, and tensile testing. Biocompatibility was assessed using primary cardiac fibroblasts, with cell viability evaluated by MTT assay and fluorescence microscopy.

## 2. Materials and Methods

### 2.1. Materials

Tecophilic™ SP-80A-150 polyurethane (PU) pellets were supplied by Lubrizol, USA. Penicillin/Streptomycin (Pen/Strep) and phosphate-buffered saline (PBS) were purchased from Wisent, Canada. Reagents, including concentrated sulfuric acid (H_2_SO_4_; 98 %), phosphoric acid (H_3_PO_4_), potassium permanganate (KMnO_4_), hydrogen peroxide (H_2_O_2_; 30%), hydrochloric acid (HCl; 37%), L-ascorbic acid, ammonia solution (25%), N,N-dimethylformamide (DMF), 3-(4,5-dimethylthiazol-2-yl)-2,5-diphenyltetrazolium bromide (MTT), and dimethyl sulfoxide (DMSO) were obtained from Sigma-Aldrich, USA. Fetal bovine serum (FBS), Dulbecco’s modified Eagle medium/nutrient mixture F12 (DMEM/F12), and Dulbecco’s phosphate-buffered saline (DPBS) were provided by Gibco, USA. Tetrahydrofuran (THF) was acquired from Acros Organics, USA. Phalloidin-iFluor 594 reagent was purchased from Abcam, UK. Live/dead assay (Calcein-AM and ethidium homodimer-1 (EthD-1)) was obtained from Invitrogen, USA. Ethylene (99.5%) and ammonia (99.99%) gases were provided by Air Liquide, Canada.

### 2.2. rGO synthesis

Reduced graphene oxide (rGO) was synthesized using our previously reported and well-characterized protocol [31]. Briefly, graphene oxide (GO) was first prepared using a modified Hummers’ method by dispersing 3 g of graphite in H_2_SO_4_/H_3_PO_4_ (360/40 mL), followed by the addition of 18 g KMnO_4_ and reaction at 50 °C for 24 h. The mixture was diluted with ice and quenched with 30% H_2_O_2_, then centrifuged and repeatedly washed with HCl and deionized (DI) water. The purified GO was dispersed in DI water. For reduction, GO (1 mg mL ¹) was mixed with ascorbic acid (10:1, w/w), adjusted to pH ∼10 with ammonia, and heated at 95 °C for 4 h. The resulting rGO was vacuum-filtered, washed to neutral pH, and dried at 60 °C overnight.

### 2.3. Electrospinning

#### 2.3.1. Solution preparation

Electrospinning solutions were prepared with or without rGO. For the control formulation, PU pellets were dissolved at 5 wt% in a 2:1 (v/v) mixture of THF and DMF, and the mixture was magnetically stirred overnight. For rGO-containing formulations, rGO was first ultrasonicated in DMF for 1 h, after which THF and PU pellets were added to obtain solutions containing 5 wt% PU and either 0.5 or 1 wt% rGO while maintaining a 2:1 THF:DMF ratio. These suspensions were stirred overnight and briefly bath-sonicated for 5 min immediately before electrospinning.

#### 2.3.2. Electrospinning parameters

The electrospinning solution was loaded into a syringe fitted with a 21G needle and placed in a Tong Li top-down electrospinning setup. The flow rate was adjusted to 1.5 mL/h, and the mandrel (10 cm in diameter) was rotated at 50 rpm to produce randomly oriented fibers (referred to as random fibers) or at 1000 rpm to produce aligned fibers. To create a mat with both random and aligned fibers, 5/6 of the solution was electrospun at 50 rpm, followed by the remaining 1/6 at 1000 rpm. The needle moved along the length of the mandrel at 10 mm/s. For solutions without rGO, the voltage was set to 14 kV, and the distance between the needle and collector was 14 cm. For solutions containing rGO, the voltage and distance were adjusted to 15 kV and 15 cm, respectively. After electrospinning, the mats were carefully removed from the aluminum foil and left to dry in ambient air before further use.

### 2.4. Plasma polymerized coating

To improve cell adhesion on the mats, a previously developed coating (LPPE:N) was applied [30]. The electrospun samples were placed in a low-pressure chamber where plasma was generated and sustained by introducing ethylene (C_2_H_4_) and ammonia (NH_3_) gases along with an RF current. The chamber pressure was maintained at approximately 600 mTorr, with ethylene and ammonia gas flow rates set at 20 and 15 sccm, respectively. The power was adjusted to 15 W, and the samples were exposed to the plasma for 15 minutes. The coated samples were used within four days of coating.

### 2.5. Scanning electron microscopy

The electrospun mats were prepared for scanning electron microscopy (SEM) by sputter-coating them under vacuum with a 20 nm thick layer of chromium. The coated samples were then mounted on a sample holder using double-sided tape and imaged at 15 kV. Fiber diameters were measured using ImageJ software (NIH). For each mat, five images were analyzed, with 20 fibers per image, for a total of 100 fibers. Fiber orientation, expressed in degrees (0 to 180), was also assessed for 100 fibers across five images per mat by tracing along the fiber’s orientation and recording its angle; horizontal and vertical lines corresponded to 0° and 90° angles, respectively.

### 2.6. X-ray photoelectron spectroscopy

X-ray photoelectron spectroscopy (XPS) analysis was carried out using a VG ESCALAB 250Xi system with a monochromatic Al Ka source operating at 14.7 kV and 14.9 mA. The analyzed area was approximately 1 mm², with a sampling depth of less than 10 nm. Spectra were collected at a 0° emission angle (perpendicular to the mat surface). Data quantification was done using Avantage v4.12 software (Thermo Fisher Scientific, USA), based on integral peak intensities and sensitivity factors from the Al Thermo table. Charging effects were corrected by referencing all peaks to the carbon (C1s) peak at a binding energy of 285.0 eV.

### 2.7. Tensile testing

Tensile testing of the electrospun mats was conducted using a Mach-1 device (Biomomentum, Canada) with a 10 N load cell and a test speed of 20 mm/min. The samples had a width of approximately 10 mm, and the gauge length between the grips was 23 mm. The thickness of each mat was measured at three different points using a micrometer and then averaged. Five samples were tested for each mat. For wet mat testing, the samples were soaked in deionized water for 24 hours prior to testing. Force and position data were recorded until the sample either broke or the axis limit (∼10 cm) was reached. Load, tensile stress, and tensile strain were calculated based on this data and the sample dimensions, and a stress-strain curve was plotted. From these curves, Young’s modulus and tensile strength were determined.

### 2.8. Electrical conductivity

Volume resistivity measurements were carried out using a Keithley 6517A resistivity meter paired with the 8009 resistivity test fixture, with samples having dimensions larger than 54 mm. The fibers were randomly oriented, and the average thicknesses of PU, PU rGO 0.5%, and PU rGO 1% mats were 400 µm, 660 µm, and 1050 µm, respectively. A voltage of 1 volt was applied to the top electrode, and the current measured at the bottom electrode was recorded. The value after 60 seconds of electrification was used, and conductivity was calculated using the following formulas:

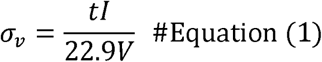

Where σ□ represents the volume conductivity (S/cm), 22.9 is the surface area of the sample under the test electrode, V is the applied voltage, t is the average sample thickness, and I is the measured current [32]. All measurements were performed on three independent replicates for each mat.

### 2.9. Cell culture

Primary cardiac fibroblasts, previously harvested from neonatal Sprague-Dawley rats (CRCHUSJ, protocol number: 2023-4833) and frozen following standard procedures [33, 34], were thawed and cultured for use in subsequent experiments. The cells were plated in a culture flask and grown in a DMEM/F12 medium supplemented with 10% FBS and 1% Pen/Strep. They were incubated in a humidified environment at 37°C with 5% CO_2_, and the medium was refreshed every 2 to 3 days until the cells reached about 80% confluence. The cells were then detached using trypsin, counted, and seeded for further experiments, with passages between 3 and 6 used for the studies.

### 2.10. Sample Sterilization

The electrospun mats were cut into discs of various diameters to fit wells of different sizes, depending on the experiment. The mats were sterilized by first wetting and washing them in an antibiotic solution containing 1% Pen/Strep in PBS. They were then exposed to UV light for at least 90 minutes.

### 2.11. Indirect cytotoxicity assessment

To evaluate the indirect cytotoxicity of the electrospun mats, cells were cultured in medium that had been in contact with the mats, in accordance with ISO 10993-5 standards [35]. In summary, the mats were cut into 7 mm diameter discs, sterilized as previously described, and placed in wells of a 48-well plate, each covered with 400 µL of medium. Wells without mats (Tissue Culture Plate, TCP) served as controls. The plates were incubated at 37°C for 4 days, after which the extracts were used to replace the medium of cells previously seeded in a 96-well plate at a density of 20,000 cells per well. After 24 hours of incubation, further assays were conducted.

### 2.12. MTT assay

The MTT assay was performed to assess cell viability for each condition based on metabolic activity [36]. This method involves the conversion of MTT into formazan crystals by living cells, which are then dissolved for optical absorbance measurement. The absorbance correlates with the number of viable cells. To prepare the MTT solution, MTT powder was vortexed in PBS at a concentration of 5 mg/mL, sterilized by filtration (pore size < 0.2 µm), and mixed with DMEM/F12 at a 1:9 ratio. 100 µL of this solution was added to each well of a 96-well plate, replacing the medium. The plate was incubated at 37°C for 4 hours. After incubation, the solution was replaced with 100 µL of DMSO to completely dissolve the formazan crystals. Optical absorbance was measured at 570 nm using a microplate reader, and cell viability for each condition was calculated by comparing the absorbance to that of the control, where viability was set at 100%.

### 2.13. Live/Dead assay

In addition, a Live/Dead assay was conducted. This method uses two dyes: one stains live cells green and the other stains dead cells red [37, 38]. First, the cell media was removed from the wells, and the cells were gently rinsed with DPBS two or three times. Next, 100 µL of the Live/Dead solution (prepared by mixing 1 µL of Calcein-AM and 2 µL of EthD-1 in 2 mL of DPBS) was added to each well, and the plate was incubated at 37°C for 30 minutes. After incubation, the solution was discarded, and the cells were washed again with DPBS two or three times before being imaged with a Leica SP8-DLS confocal microscope (Leica Biosystems, Germany).

### 2.14. Statistical analysis

Statistical analyses were carried out using GraphPad Prism software. To assess the significance of differences between groups, a one-way analysis of variance (ANOVA) was conducted, followed by Tukey’s multiple comparisons test. Data are shown as mean ± standard deviation where appropriate, and the number of replicates (n) for each experiment is indicated.

## 3. Results and Discussion

This study focused on developing a biocompatible and conductive electrospun patch for cardiac tissue regeneration. It explored material selection, fabrication techniques, characterization, and in vitro testing. PU was chosen as the primary material for its mechanical properties, elasticity, and biocompatibility, closely matching native myocardial tissue, while rGO was incorporated to enhance electrical conductivity. Five different electrospun mats were fabricated and are summarized in **Table 1** according to their composition and fiber orientation. These included randomly oriented PU fibers (PU R), fully aligned PU fibers (PU A), a bilayer mat with a random base and aligned top layer (PU R-A), and two conductive bilayer variants containing 0.5% and 1% rGO (PU rGO0.5 and PU rGO1, respectively).

**Table 1.** Description of fabricated electrospun mats.

| **Name** | **Composition** | **Fiber orientation** |
|---|---|---|
| PU R | Tecophilic™ 5% | Random |
| PU A | Tecophilic™ 5% | Aligned |
| PU R-A | Tecophilic™ 5% | Random – Aligned |
| PU rGO0.5 | Tecophilic™ 5% rGO 0.5% | Random – Aligned |
| PU rGO1 | Tecophilic™ 5% rGO 1% | Random – Aligned |

### 3.1. Fiber Morphology

Electrospinning produced continuous, highly porous nanofibrous mats for all formulations, closely resembling the fibrillar architecture of native extracellular matrix (**Fig. 2**). Neat PU mats exhibited smooth, uniform fibers arranged either randomly (PU R), in a uniformly aligned monolayer (PU A), or in a bilayer random–aligned architecture (PU R-A) (**Fig. 2a-d**). At higher collector rotation speeds used to generate aligned fibers, slight fiber coiling was observed, most prominently in PU A and PU R-A samples (**Fig. 2b-c**). This phenomenon could be attributed to axial shrinkage of the deposited mat upon removal from the collector, which induces compressive buckling of the fibers. Interestingly, such micro-scale waviness has been reported to be beneficial for cardiac cell function, as coiled fibers can deform with contracting cells, reducing mechanical resistance and enabling greater contractile amplitude and synchronization compared to straight fibers [39]. Therefore, the observed morphology in aligned and random-aligned PU mats could be advantageous for cardiac tissue engineering.

**Fig. 2.**
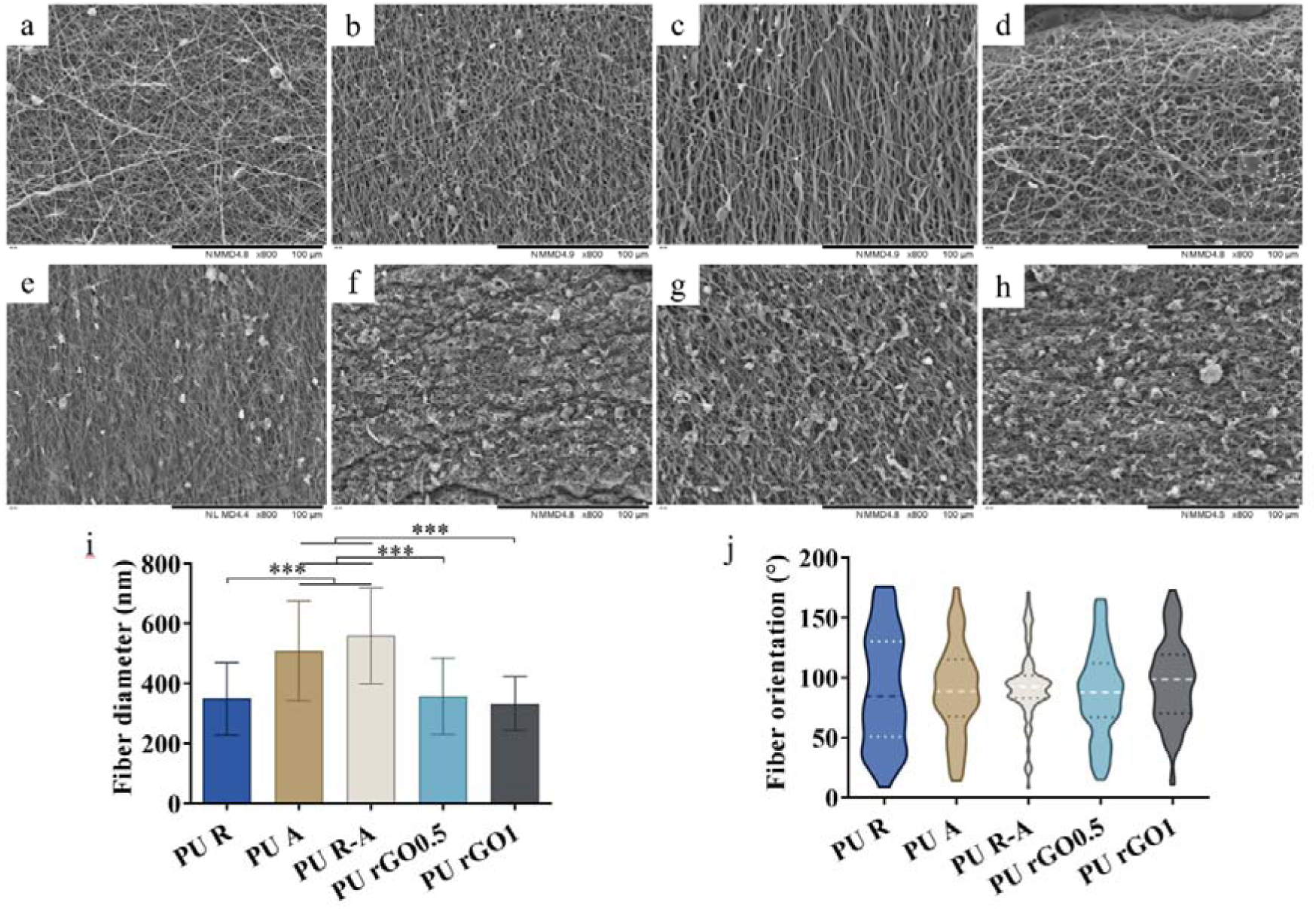
SEM micrographs and quantitative analysis of electrospun PU mats with and without rGO. Representative SEM images show (a–d) PU mats without rGO and (e–h) PU mats containing rGO. (a) PU R, (b) PU A, (c) PU R-A (front side), (d) PU R-A (back side), (e) PU rGO0.5 (front side), (f) PU rGO0.5 (back side), (g) PU rGO1 (front side), and (h) PU rGO1 (back side). (i) Average fiber diameters of the electrospun mats (n = 100). (j) Fiber orientation distributions for the corresponding mats are shown as violin plots, with median and interquartile ranges (n = 100). *** indicates statistically significant differences of p < 0.001.

The incorporation of rGO markedly altered fiber surface morphology. Both PU-rGO0.5 and PU-rGO1 mats exhibited increased surface roughness and frequent bead-like features along the fibers (**Fig. 2e–h**), arising from local agglomeration of rGO sheets within the polymer jet during electrospinning. These nanoscale protrusions increase the available surface area and introduce topographical cues that are known to promote protein adsorption and integrin-mediated cell attachment, which are critical for cardiac cell adhesion and maturation.

### 3.2. Fiber Diameter

Quantitative analysis revealed that fiber diameter was strongly influenced by both collector speed and rGO incorporation (**Fig. 2i and S1**). Random PU fibers (PU R) displayed an average diameter of 349 ± 120 nm, while aligned PU mats showed significantly thicker fibers, with diameters of 508 ± 167 nm for PU A and 559 ± 160 nm for PU R-A. This increase could be attributed to the reduced jet stretching time caused by rapid fiber deposition on the rotating drum, which limits solvent evaporation and elongational thinning. The random-aligned hybrid structure (PU R-A) exhibited the largest diameter, likely due to the mechanical constraint imposed by the underlying random layer, which partially restricts further fiber drawing during deposition of the aligned layer.

In contrast, incorporation of rGO produced a clear reduction in fiber diameter, yielding 357 ± 127 nm for PU-rGO0.5 and 333 ± 90 nm for PU-rGO1. This effect is consistent with the increased electrical conductivity of the spinning solution, which enhances jet stretching through stronger electrostatic repulsion forces. Notably, all mats exhibited fiber diameters within the 300–600 nm range, closely matching the scale of collagen and elastin fibers in native myocardial extracellular matrix [40]. This biomimetic size range is known to promote cardiomyocyte alignment, sarcomere organization, and mechano-electrical coupling, reinforcing the suitability of these scaffolds for cardiac applications [41].

### 3.3. Fiber Orientation

Fiber alignment was quantified using orientation analysis and visualized using violin plots (**Fig. 2j and S2**). As expected, PU R mats exhibited a uniform distribution of orientations, confirming their random architecture. In contrast, PU A and PU R-A mats showed pronounced peaks centered near 90°, indicating strong uniaxial alignment. The PU R-A structure displayed the tightest distribution, with 50% of fibers oriented between 83° and 102°, reflecting superior alignment compared to PU A. This enhanced organization is attributed to the mechanically stabilizing random base layer, which limits post-deposition shrinkage and preserves fiber straightness in the aligned top layer.

The addition of rGO slightly broadened the orientation distributions, indicating a modest reduction in alignment relative to PU R-A. This likely arises from increased jet instability caused by heterogeneous electrical conductivity and rGO agglomerates [42, 43]. Nevertheless, both PU-rGO0.5 and PU-rGO1 mats remained substantially more aligned than PU R, retaining a preferential fiber orientation that is essential for guiding cardiomyocyte elongation and anisotropic electrical conduction.

Importantly, the degree of alignment achieved in PU R-A and rGO-containing mats closely mirrors the anisotropic architecture of native myocardium, where aligned collagen fibers and cardiac cells enable directional force transmission and rapid electrical propagation. Thus, the structural features of these electrospun patches are well-suited to support functional cardiac tissue formation.

These characterizations emphasize how both fiber orientation and rGO incorporation affect the structural properties of the electrospun mats. The aligned PU R-A mats, along with the increased roughness in PU rGO0.5 and PU rGO1 mats, indicate that combining a random-aligned fiber architecture with rGO incorporation enables fine control over fiber diameter, surface topography, and anisotropy. The PU R-A platform provides the highest degree of alignment and mechanical stability, while rGO introduces nanoscale roughness without disrupting the biomimetic fiber size range. This modification could potentially influence cell adhesion and the mechanical properties of the mats.

### 3.4. Mechanical Properties

The uniaxial tensile behavior of the electrospun mats revealed characteristic nonlinear stress–strain profiles for all samples except PU R, exhibiting two distinct quasi-linear regions (**Fig. 3a**). These regions correspond to an initial low-strain regime dominated by fiber straightening and reorientation (E1), followed by a higher-strain regime governed by fiber stretching and load transfer within the fibrous network (E2). This two-stage mechanical response closely resembles that of native soft tissues, such as myocardium, in which collagen fiber recruitment produces a strain-dependent increase in stiffness [44, 45].

**Fig. 3.**
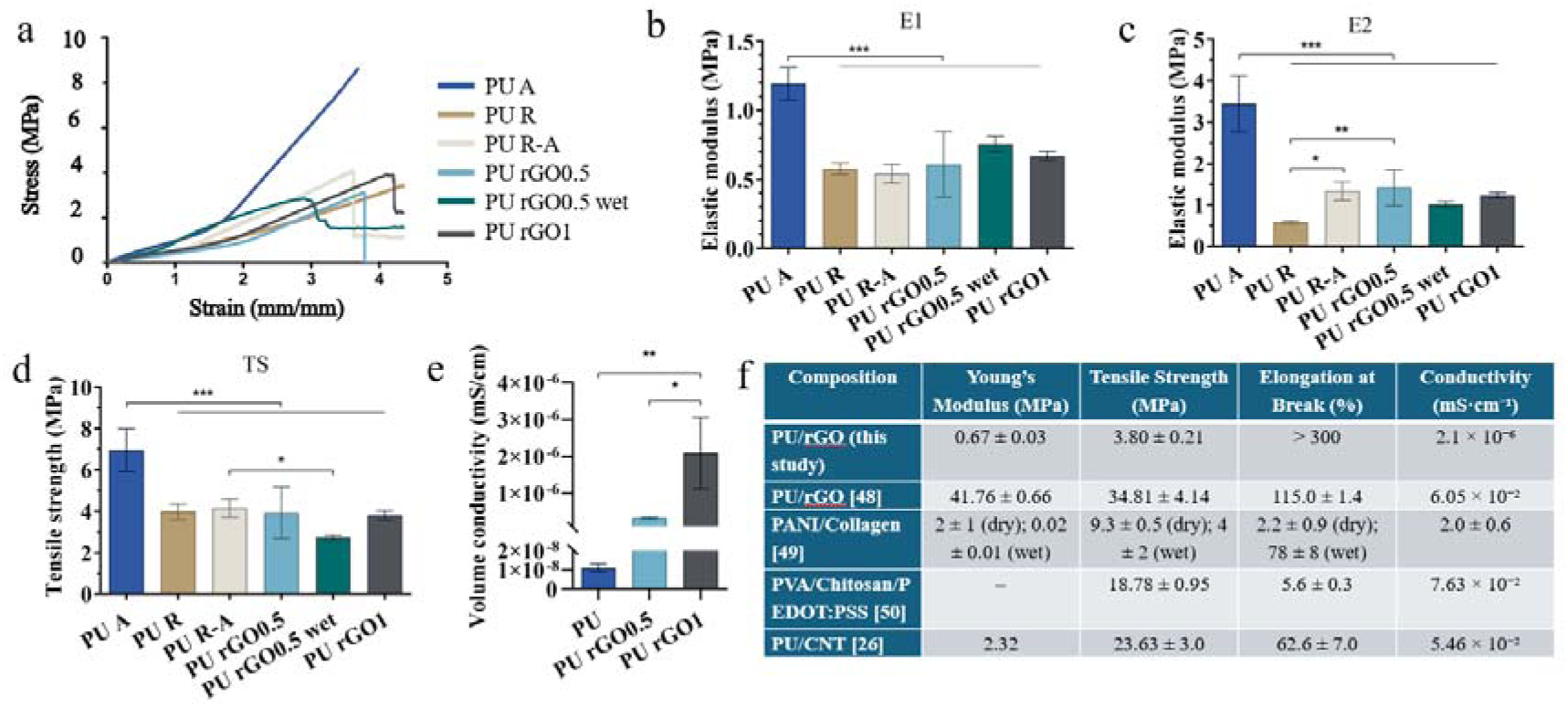
Mechanical and electrical characterization of electrospun polyurethane (PU) and PU/rGO composite mats. (a) Typical stress–strain curves obtained from uniaxial tensile testing. (b) First elastic modulus (E□) and (c) second elastic modulus (E□) derived from the initial and secondary linear regions of the stress-strain curves (n = 5). (d) Tensile strength (TS) of the electrospun mats (n = 5). (e) Volume conductivity of PU and PU/rGO composites (n = 3). Conductivity values are reported in mS·cm ¹. (f) Summary of the mechanical and electrical properties of electrospun cardiac patches from this study compared with literature data. *, **, *** indicate statistically significant differences of p < 0.05, p < 0.01, and p < 0.001, respectively.

Aligned PU mats (PU A) exhibited the highest stiffness and strength among all groups, with E1 = 1.19 ± 0.12 MPa, E2 = 3.44 ± 0.68 MPa, and tensile strength (TS) = 6.96 ± 1.04 MPa (**Fig. 3b-d**). This superior mechanical performance could be attributed to the strong fiber alignment parallel to the loading direction, which enables efficient stress transfer along continuous polymer chains and minimizes fiber slippage. These results confirm that fiber orientation is a dominant determinant of mechanical anisotropy in electrospun PU scaffolds.

In contrast, both random PU (PU R) and random-aligned PU (PU R-A) exhibited significantly lower stiffness and tensile strength. PU R-A showed E1 = 0.54 ± 0.07 MPa, E2 = 1.34 ± 0.23 MPa, and TS = 4.16 ± 0.44 MPa, values that were in the range of those measured for PU R (E1 = 0.58 ± 0.04 MPa, E2 = 0.58 ± 0.04 MPa, TS = 3.98 ± 0.37 MPa). This indicates that the random fiber base layer in PU R-A partially offsets the mechanical reinforcement provided by the aligned top layer. Importantly, this mechanical moderation brings PU R-A into a stiffness range closer to native myocardium, which exhibits end-diastolic moduli of approximately 0.2-0.5 MPa and relatively low tensile strength. Thus, the bilayer random-aligned architecture provides a favorable compromise between mechanical compliance and surface anisotropy for cardiac cell alignment.

Furthermore, incorporation of rGO did not significantly alter the tensile mechanics of the mats. Both PU-rGO0.5 and PU-rGO1 showed elastic moduli and tensile strengths comparable to PU R-A (**Fig. 3b-d**), indicating that rGO at these concentrations does not compromise the mechanical integrity of the PU fiber network. This is advantageous for cardiac patch applications, as electrical functionality can be introduced without stiffening the scaffold.

On the other hand, hydration produced a modest but measurable softening effect. Compared with dry PU-rGO0.5 mats, wet PU-rGO0.5 showed reduced tensile strength (from 3.92 ± 1.25 MPa to 2.74 ± 0.11 MPa) and lower E2 (from 1.42 ± 0.44 MPa to 1.01 ± 0.07 MPa), consistent with water-induced disruption of intermolecular hydrogen bonding and partial plasticization of the PU matrix. Nevertheless, the mats maintained sufficient structural integrity under wet conditions relevant to physiological environments.

Moreover, all mats exhibited elongation at break exceeding 300%, reflecting the intrinsic elasticity of PU and the highly deformable fibrous architecture. This extensibility far exceeds the strain experienced by myocardium during the cardiac cycle (∼20–30%), ensuring that the patches can accommodate repeated physiological deformation without mechanical failure.

Overall, these results demonstrate that the bilayer PU R-A and PU-rGO mats achieve a mechanically compliant yet resilient profile that closely matches cardiac tissue requirements, while preserving surface fiber alignment for cardiac applications.

### 3.5. Electrical Conductivity

The electrical conductivity of the electrospun mats increased significantly with rGO incorporation (**Fig. 3e**). Pure PU mats exhibited very low conductivity (1.1 ± 0.2 × 10^-8^ mS/cm), reflecting the intrinsically insulating nature of polyurethane. Incorporation of 0.5 wt% rGO increased conductivity by nearly one order of magnitude to 3.2 ± 0.3 × 10^-7^ mS/cm, while 1 wt% rGO produced a further significant increase to 2.1 ± 0.9 × 10^-6^ mS/cm. These results indicate the formation of a progressively more connected conductive network within the PU fiber matrix as rGO content increases, consistent with percolation-type behavior observed in other polymer-graphene composites [46, 47].

Although these values remain substantially lower than the effective electrical conductivities of native myocardium (∼2.1 mS/cm longitudinally and 0.5 mS/cm transversely), the dry-state measurements likely underestimate *in vivo* performance. In physiological conditions, the presence of ion-rich extracellular fluid (∼20 mS/cm) is expected to significantly enhance effective electrical coupling across the scaffold. Moreover, even modest increases in scaffold conductivity have been shown to improve intercellular electrical communication, calcium wave propagation, and synchronized beating in cardiac constructs, suggesting that the rGO-based mats developed here provide a functionally relevant level of electrical enhancement.

### 3.6. Electromechanical Performance and Biomimicry

A key advantage of the PU/rGO electrospun scaffolds lies in their unique combination of high deformability, low stiffness, and enhanced electrical conductivity. As summarized in **Fig. 3f**, the PU/rGO mats from this study exhibit one of the lowest Young’s moduli (0.67 ± 0.03 MPa) and tensile strengths (3.80 ± 0.21 MPa) reported for conductive electrospun cardiac patches, while maintaining an elongation at break exceeding 300%. This mechanical profile is well-matched to the highly dynamic and compliant nature of myocardial tissue, which undergoes repeated 20-30% cyclic strain during normal cardiac function. Compared with previously reported conductive scaffolds such as PU/rGO [48], PANi/collagen [49], PVA/chitosan/PEDOT:PSS [50], and PU/CNT [26], the present PU/rGO patches are far more stretchable but substantially softer. While this leads to lower ultimate strength, it greatly reduces mechanical mismatch with the heart, which is known to contribute to inflammation, fibrosis, and impaired electromechanical integration. Therefore, the combination of moderate stiffness, high extensibility, and suitable conductivity suggests superior long-term electromechanical compatibility for epicardial or myocardial patch applications.

### 3.7. Chemical composition of the plasma-polymerized coating

XPS analysis was used to evaluate the surface chemistry of electrospun PU/rGO fibers before and after deposition of the LPPE:N coating. **Fig. 4** shows high-resolution C1s, O1s, and N1s spectra for uncoated and coated mats, allowing direct comparison of chemical functionalities introduced by the plasma process.

**Fig. 4.**
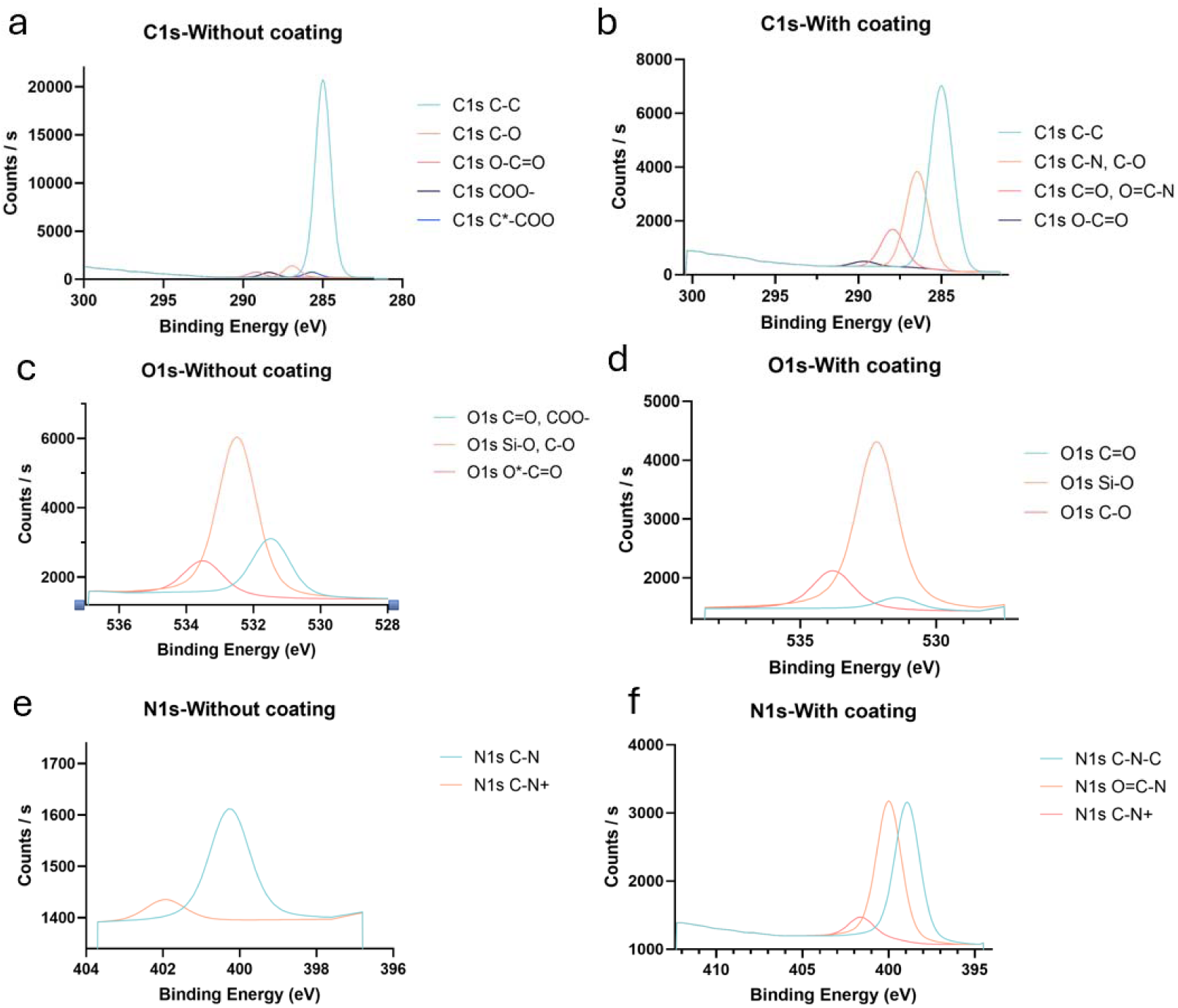
High-resolution XPS spectra of electrospun PU/rGO0.5 fibers before and after LPPE:N plasma coating. (a,b) C1s spectra of uncoated and coated fibers, showing the emergence of C–N and increased polar functionalities after coating. (c,d) O1s spectra revealing enhanced C–O and C=O environments following plasma polymerization. (e,f) N1s spectra demonstrating a dramatic increase in nitrogen-containing groups (C–N–C, O=C–N, and C–N□) after coating, confirming successful deposition of a nitrogen-rich plasma polymer layer.

#### 3.7.1. Carbon chemistry (C1s)

The uncoated fibers (**Fig. 4a**) display a dominant C–C/C–H peak at ∼284.8 eV, characteristic of the PU backbone, along with weaker contributions from C–O (∼286.2 eV) and O–C=O (∼288.5–289.0 eV) associated with urethane and ester groups. After plasma coating (**Fig. 4b**), the C1s envelope becomes more complex, showing clear contributions from C–N and C–O bonds (∼286 eV), as well as C=O and O–C=O species. The appearance of a strong C–N component confirms the incorporation of nitrogen-containing organic groups into the surface layer. These changes indicate that the original PU surface was effectively covered by a thin plasma-polymerized film enriched with polar and nitrogen-bearing functionalities.

#### 3.7.2. Oxygen chemistry (O1s)

The O1s spectrum of the uncoated fibers (**Fig. 4c**) is dominated by components attributed to C=O/COO–(∼531.5 eV) and Si–O/C–O (∼532.8–533.5 eV), arising from the polymer and residual processing components. After coating (**Fig. 4d**), the O1s profile changes markedly, with an increased contribution of C–O and C=O environments and the appearance of Si–O species, consistent with the formation of a plasma-derived organic network. These oxygen-containing functional groups increase surface polarity and wettability, which are known to promote protein adsorption and subsequent cell adhesion.

#### 3.7.3. Nitrogen chemistry (N1s)

The most significant change is observed in the N1s region. The uncoated PU/rGO0.5 fibers (**Fig. 4e**) show only a weak nitrogen signal (∼0.7 at.%), mainly associated with intrinsic urethane C–N bonds and trace charged amines. In contrast, the coated fibers (**Fig. 4f**) exhibit a strong N1s signal with multiple components corresponding to C–N–C (∼399.5 eV), O=C–N (∼400.5 eV), and protonated or positively charged amines (C–N□, ∼401.5–402 eV). Quantitatively, the nitrogen content increases dramatically to 17.8 at.%, confirming the successful deposition of the nitrogen-rich plasma polymer layer.

The observed nitrogen level is consistent with previous reports for LPPE:N coatings applied to electrospun PET fibers, where nitrogen contents of ∼13% were achieved and correlated with improved cell adhesion [29]. The presence of primary and secondary amine groups is particularly important, as these functionalities provide positive surface charge and reactive sites for protein binding, which enhances integrin-mediated cell attachment and spreading [51].

The combined enrichment in nitrogen-containing groups and polar oxygen functionalities demonstrates that the LPPE:N coating substantially alters the surface chemistry of the electrospun fibers. Such surfaces are well known to promote adsorption of ECM proteins in favorable conformations, thereby improving cell adhesion and survival. Preliminary biological testing in this study already indicated improved cardiac cell viability on coated mats, although further quantitative analysis of amine density and long-term cell adhesion is still required [28]. Beyond nitrogen functionalization, plasma-polymerized coatings can also be tuned to incorporate oxygen-rich or bioactive moieties. Previous studies have shown that oxygen-functional plasma coatings can enhance muscle cell alignment and contractility [52], suggesting that similar strategies could be used to further optimize cardiac performance on heart-on-a-chip platforms. Overall, the XPS results confirm that LPPE:N provides a robust and chemically active interface that is highly suitable for improving cell-scaffold interactions in cardiac patches.

### 3.8. *In vitro* Biocompatibility

The indirect cytotoxicity of electrospun PU/rGO mats, with and without LPPE:N surface modification, was evaluated using MTT metabolic activity and Live/Dead fluorescence assays. This extract-based approach probes whether any leachable species from the materials adversely affect cell viability, providing a stringent assessment of biocompatibility for tissue engineering applications.

#### 3.8.1. MTT assay

As shown in **Fig. 5a**, all material extracts exhibited cell viabilities above the TCP control (100%), confirming the absence of indirect cytotoxicity. Cells exposed to the PU R-A extract displayed the highest metabolic activity (128 ± 15%), followed closely by PU rGO0.5 (123 ± 17%). These values suggest that the base PU scaffold and low rGO loading do not release harmful by-products and may even promote cellular metabolic activity, likely due to favorable surface chemistry and low oxidative stress.

**Fig. 5.**
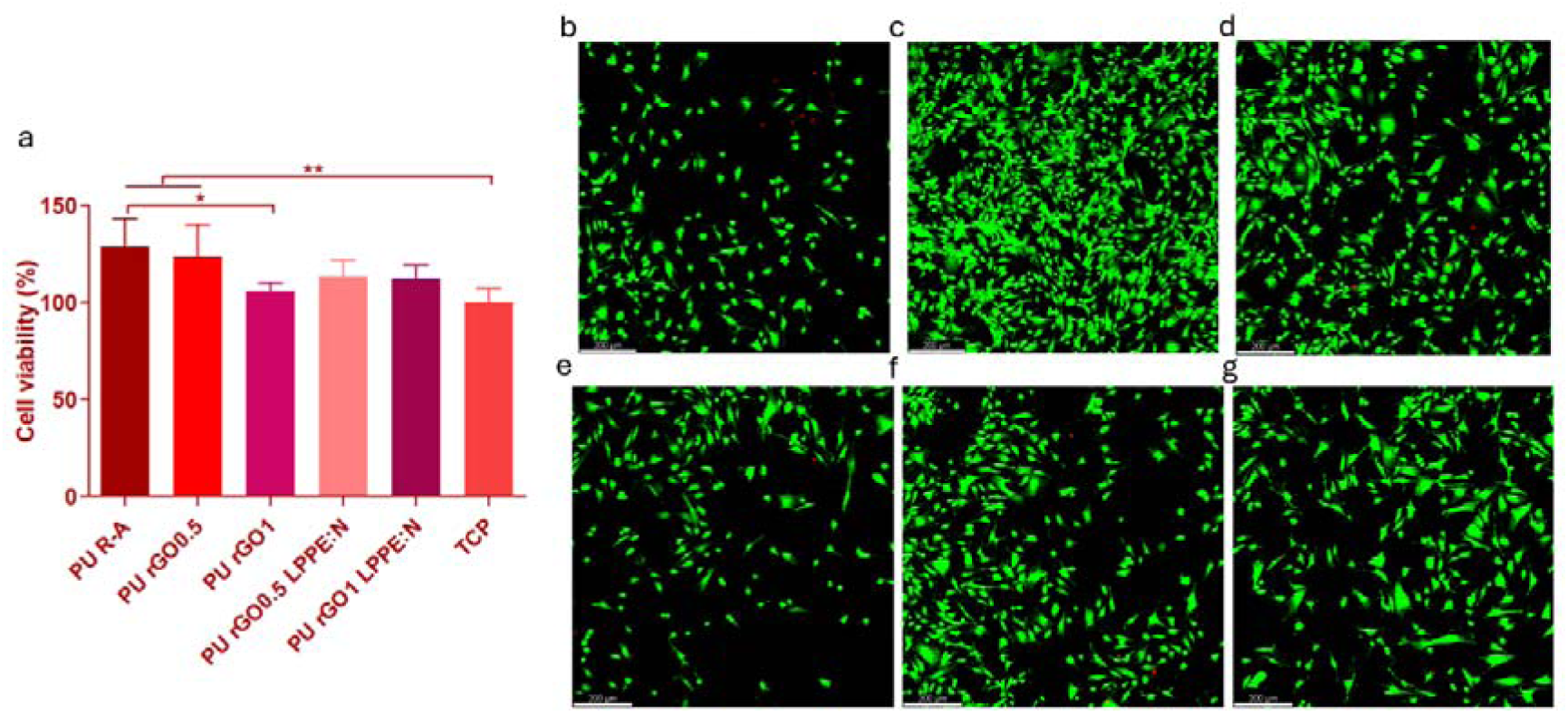
(a) Cell viability evaluated via MTT reduction in indirect cytotoxicity assessment of electrospun mats, normalized to TCP control. (n=6). Live/Dead staining images resulting from indirect cytotoxicity assessment of different electrospun mats for (b) TCP control, (c) PU R-A mat, (d) PU rGO0.5 mat, (e) PU rGO1 mat, (f) PU rGO0.5 LPPE:N mat, and (g) PU rGO1 LPPE:N mat (n=6). * and ** indicate statistically significant differences of p < 0.05 and p < 0.01, respectively.

In contrast, PU rGO1 induced a statistically significant reduction in metabolic activity compared to PU R-A (105 ± 6%), although viability remained comparable to TCP group. This decrease in viability at higher rGO concentrations is consistent with previous reports showing that higher graphene-based filler content can increase oxidative stress or disrupt ionic homeostasis by releasing adverse by-products, resulting in dose-dependent cytotoxic effects on cultured cells. Importantly, application of the nitrogen-rich plasma coating mitigated this effect. Both PU rGO0.5 LPPE:N and PU rGO1 LPPE:N exhibited increased viability compared to uncoated PU rGO1. This improvement indicates that the LPPE:N coating acts as a stabilizing and bioactive interface, likely reducing exposure to rGO-related leachables while introducing amine-rich chemistry that supports cellular metabolism.

It should be noted that MTT measures mitochondrial activity rather than direct cell number; thus, elevated values may reflect increased metabolic demand or mild stress rather than proliferation [53]. Nevertheless, the data robustly demonstrate that none of the materials induce cytotoxic leaching compared to TCP control.

#### 3.8.2. Live/Dead assay

Live/Dead staining further confirmed the MTT results. In all conditions, more than 90% of cells stained positive for calcein (green), indicating intact membranes and metabolic activity, while only a few ethidium homodimer-positive (red) cells were observed. Cells maintained a spindle-like morphology typical of healthy cardiac fibroblasts, indicating preservation of their native phenotype [54].

TCP control cultures (**Fig. 5b**) showed normal cell density and morphology. A notably higher cell density was observed in the PU R-A extract (**Fig. 5c**), consistent with its elevated MTT signal. PU rGO0.5 extracts also supported high cell numbers (**Fig. 5d**), whereas PU rGO1 extracts resulted in visibly fewer cells (**Fig. 5e**), corroborating the reduced metabolic activity measured by MTT. Furthermore, extracts from LPPE:N-coated mats (**Fig. 5f-g**) restored cell density to levels comparable to PU rGO0.5, indicating that plasma polymerization counteracts the adverse effects associated with higher rGO content. These findings strongly support the role of nitrogen-rich surface chemistry in improving the biological response to composite electrospun scaffolds.

Together, the MTT and Live/Dead data demonstrate that PU/rGO electrospun mats are non-cytotoxic and support high cardiac fibroblast viability. The LPPE:N coating further improves biocompatibility by providing a chemically active, amine-rich interface that enhances protein adsorption and stabilizes cell-material interactions. Although these *in vitro* assays confirm good short-term biocompatibility and support cell viability, they do not fully predict long-term tissue integration or functional performance. Future studies should therefore evaluate cardiomyocyte contractility, electrical coupling, inflammatory response, and in vivo integration of the patches. Overall, these results establish PU/rGO-based electrospun patches, especially when functionalized with LPPE:N, as promising biomaterials for cardiac tissue engineering and regenerative medicine, while also highlighting opportunities for future optimization of their biointerface and functional performance.

## 4. Conclusion

This study successfully developed a biocompatible and conductive electrospun patch composed of PU and rGO for potential cardiac tissue engineering applications. The electroconductive patches demonstrated a biomimetic fibrous microstructure, tunable anisotropy, and nonlinear mechanical behavior closely resembling native myocardium, while maintaining high elasticity and compliance under physiologically relevant conditions. Incorporation of rGO significantly enhanced electrical conductivity without compromising mechanical integrity, enabling improved electromechanical compatibility compared with previously reported conductive cardiac scaffolds. Furthermore, nitrogen-rich plasma-polymerized surface functionalization introduced amine- and oxygen-containing polar groups, substantially increasing surface bioactivity. This chemical modification supported cardiac fibroblast viability and mitigated potential adverse effects associated with higher rGO loadings. While these findings highlight the potential of PU/rGO electrospun patches as a promising platform for cardiac tissue engineering, further investigations are required to assess long-term cardiomyocyte attachment, electrical coupling, and synchronous contractile behavior, as well as to evaluate inflammatory response, electrical integration, and functional performance in relevant *in vivo* models. Collectively, the simultaneous control over scaffold mechanics, anisotropic architecture, electrical conductivity, and surface bioactivity positions this platform as a versatile and translatable candidate for next-generation cardiac patches.

## Acknowledgment

Our work is supported by the Natural Sciences and Engineering Research Council of Canada (NSERC) (NSERC, RGPIN-2021-03960, DGECR-2021-00337), Fonds de Recherche du Québec Santé (FRQS) (Chercheurs-boursiers J1 (313837)), Establishment of Young Investigators (324277), Montréal TransMedTech Institute (iTMT), CRCHU Sainte-Justine (CRCHUSJ), University of Montréal and Polytechnique Montreal. A.M. gratefully acknowledges the FRQS Doctoral Scholarship, the Merit Scholarship from the Faculty of Medicine of the University of Montréal, and the Michel Bergeron Scholarship from the Department of Pharmacology and Physiology of the University of Montréal.

