## Supplemental Files for "Plasma-Polymerized Polyurethane/Reduced Graphene Oxide Electrospun Patch for Cardiac Tissue Engineering"

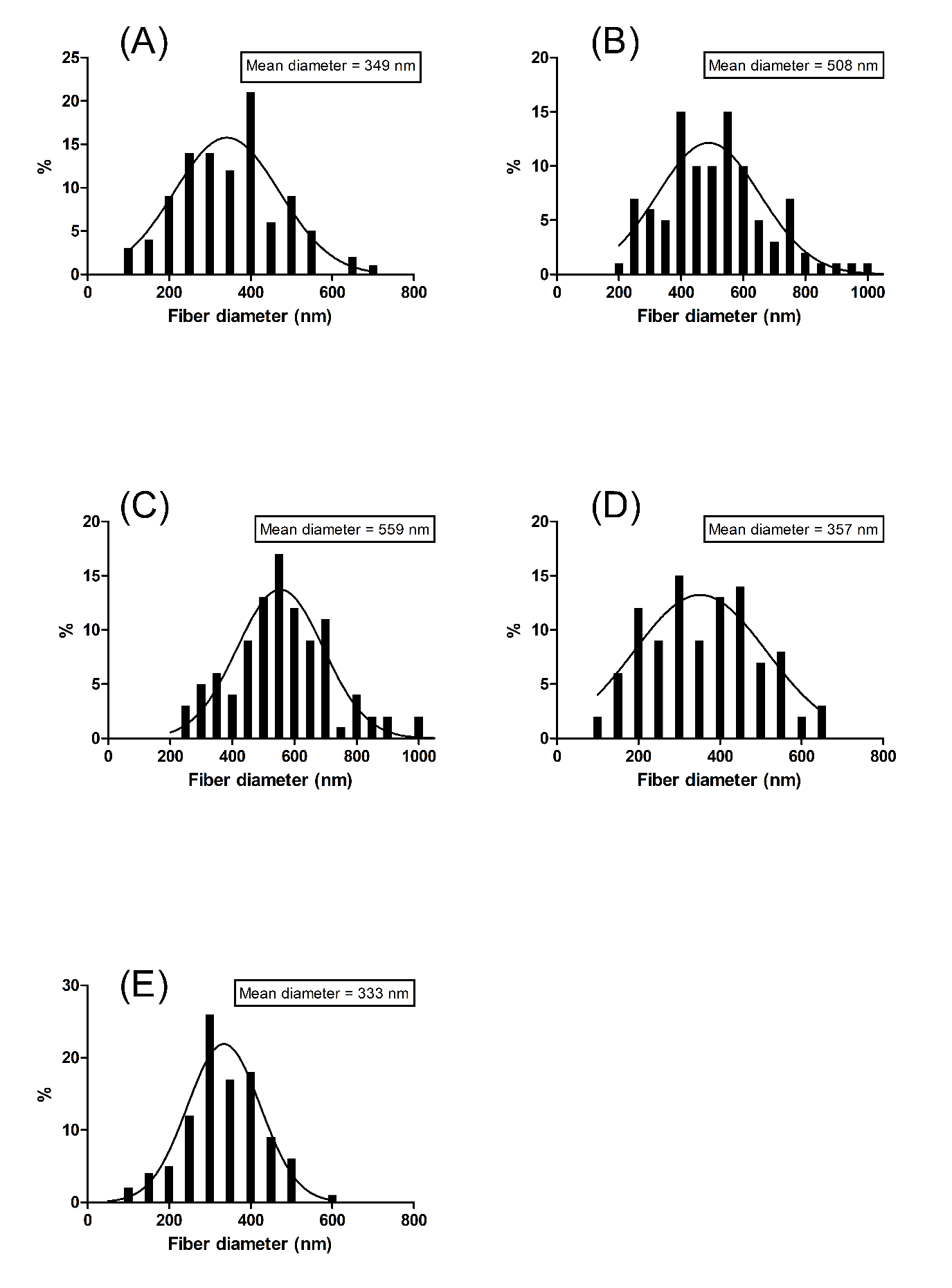
**Supplementary** **Figures**

**Fig. S1.** Fiber diameter distribution histograms of electrospun mats. (A) PU R. (B) PU A. (C) PU R-A. (D) PU rGO0.5. (E) PU rGO1. (n=100)

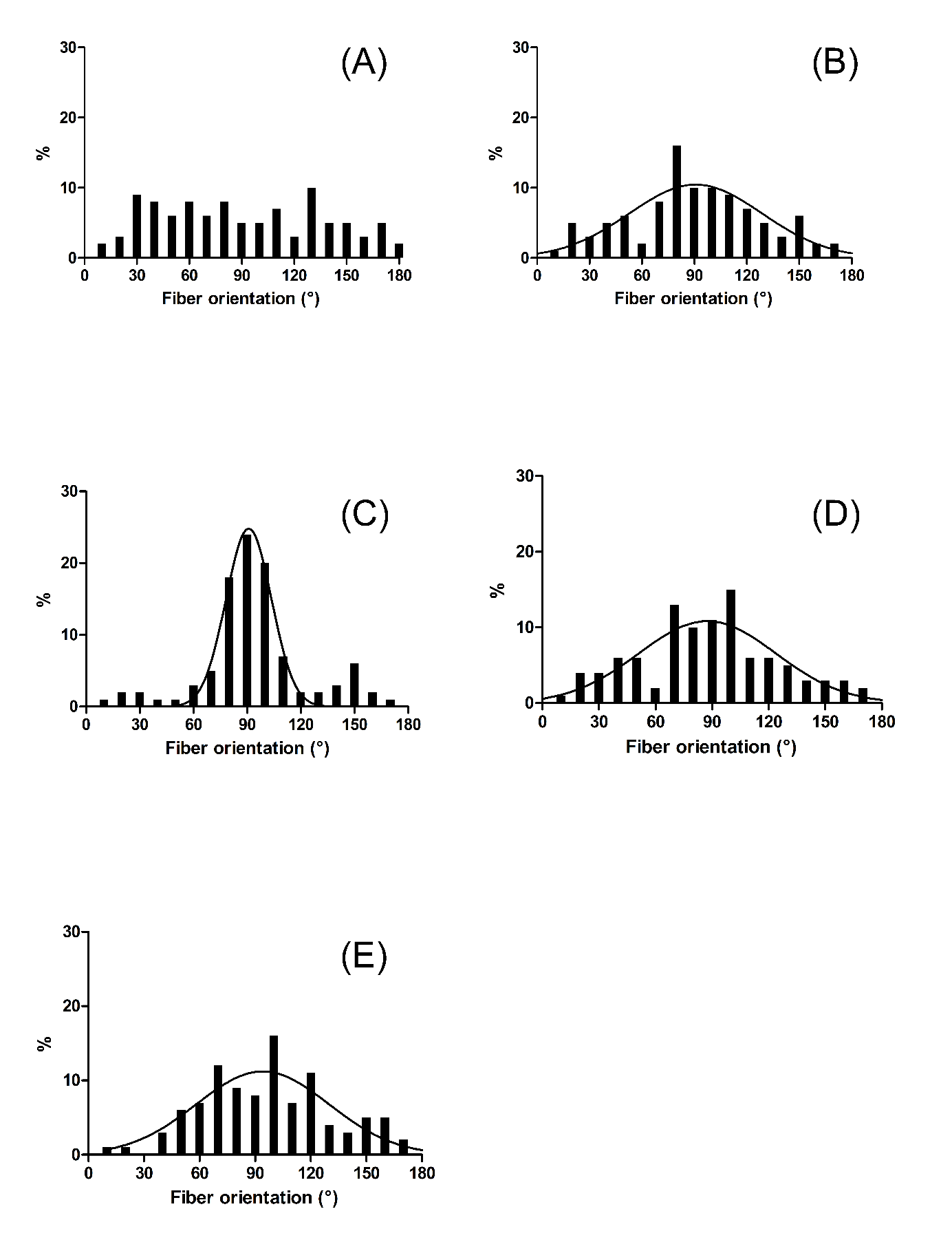

**Fig. S2.** Fiber orientation distribution histograms of electrospun mats. (A) PU R. (B) PU A. (C) PU R-A. (D) PU rGO0.5. (E) PU rGO1. (n=100)
